# scHLAcount: Allele-specific HLA expression from single-cell gene expression data

**DOI:** 10.1101/750612

**Authors:** Charlotte A. Darby, Michael J. T. Stubbington, Patrick J. Marks, Álvaro Martínez Barrio, Ian T. Fiddes

## Abstract

Studies in bulk RNA sequencing data suggest cell-type and allele-specific expression of the human leukocyte antigen (HLA) genes. These loci are extremely diverse and they function as part of the major histocompatibility complex (MHC) which is responsible for antigen presentation. Mutation and or misregulation of expression of HLA genes has implications in diseases, especially cancer. Immune responses to tumor cells can be evaded through HLA loss of function. However, bulk RNA-seq does not fully disentangle cell type specificity and allelic expression. Here we present scHLAcount, a workflow for computing allele-specific molecule counts of the HLA genes in single cells an individualized reference. We demonstrate that scHLAcount can be used to find cell-type specific allelic expression of HLA genes in blood cells, and detect different allelic expression patterns between tumor and normal cells in patient biopsies. scHLAcount is available at https://github.com/10XGenomics/scHLAcount.

## Introduction

The major histocompatibility complex (MHC) locus of human chromosome 6 is important for antigen presentation, containing genes for both class I and class II human leukocyte antigen (HLA). This locus is highly variable in the human population, with hundreds of characterized alleles that can be considerably divergent. Class I HLA alleles are responsible for neoantigen presentation, and therefore HLA haplotype information for a patient is important for developing targeted immunotherapies. Loss of HLA expression or function is likely a major driver of immunotherapy evasion. Loss of HLA class I expression has been demonstrated in relapse after immunotherapy treatment of Merkel cell carcinoma (Paulson et al. 2018) and loss of HLA class II expression was observed in relapse after hematopoietic stem-cell transplantation for acute myeloid leukemia (AML) (Christopher et al. 2018). Genomic loss of heterozygosity of HLA has been detected in 40% of non-small-cell lung cancers using the LOHHLA algorithm, which uses information about the individual’s HLA genotype to determine copy number (McGranahan et al. 2017).

In bulk RNA-seq data, expression of MHC locus genes are often underestimated due to poor mappability caused by variability in the locus. Tools that build custom diploid references such as AltHapAlignR (W. Lee et al. 2018) improves expression quantification. HLApers extended the diploid reference model to improve allele-specific expression estimates (Aguiar et al. 2019) for eQTL mapping.

We seek to apply this concept to single cell gene expression data, such as those produced by 10x Genomics’ Single Cell Immune Profiling (5’ capture) and Gene Expression (GEX) (3’ capture) Solutions. Single cell expression analysis software, such as 10x Genomics’ Cell Ranger, produce a matrix of molecule counts for each gene in each cell. HLA expression is systematically underestimated when using the reference genome compared to a personalized diploid reference (Aguiar et al. 2019). Therefore, as Cell Ranger relies on alignment to the reference genome, per-cell molecule counts for HLA genes are also likely to be underestimated, and potentially skewed by haplotype and population of origin.

HLA allele-specific expression (ASE) has been seen in lymphoblastoid cell lines (Aguiar et al. 2019). In a study of allele-specific expression of HLA-A, -B, and -C genes in peripheral blood mononuclear cells (PBMCs) subsets, no cell type specific allele preference was found (Greene et al. 2011). However, alleles in the rhesus macaque with significant cell type specific expression were found. Some HLA-C alleles with consistently higher expression have been found by qPCR (Bettens et al. 2014); this has also been observed for some alleles of the class II genes HLA-DQB1 and HLA-DQA1 (Zajacova et al. 2018).

The recent paradigm shift in solid tumor treatment by immune-checkpoint blockade (ICB) therapies has not been followed with a parallel development in prognostic biomarkers. Currently, the only FDA-approved biomarker is high PD-L1 expression (Eisenstein 2017) although many others are being investigated (Conway et al. 2018). PD-1 blockade is effective when antigens are presented by MHC of tumor cells, lymphocytes successfully infiltrate the tumor and recognize those antigens. Increased heterozygosity at HLA class I loci has shown overall better survival in ICB, especially when associated with certain HLA types (Chowell et al. 2018). HLA class II genes are also expressed in some tumor cells showing positive ICB response (Johnson et al. 2016). Here we provide a tool to study allele-specific expression of HLA genes at the single cell resolution.

## Results

scHLAcount is a postprocessing workflow for single cell gene expression data that performs allele-specific molecule counting for the main HLA class I and class II genes in each cell based on user-supplied HLA genotypes. Each molecule is assigned to an allele based on the consensus of pseudoalignments of the constituent reads to a personalized HLA reference graph. The workflow is illustrated in Figure 1.

**Figure 1:**
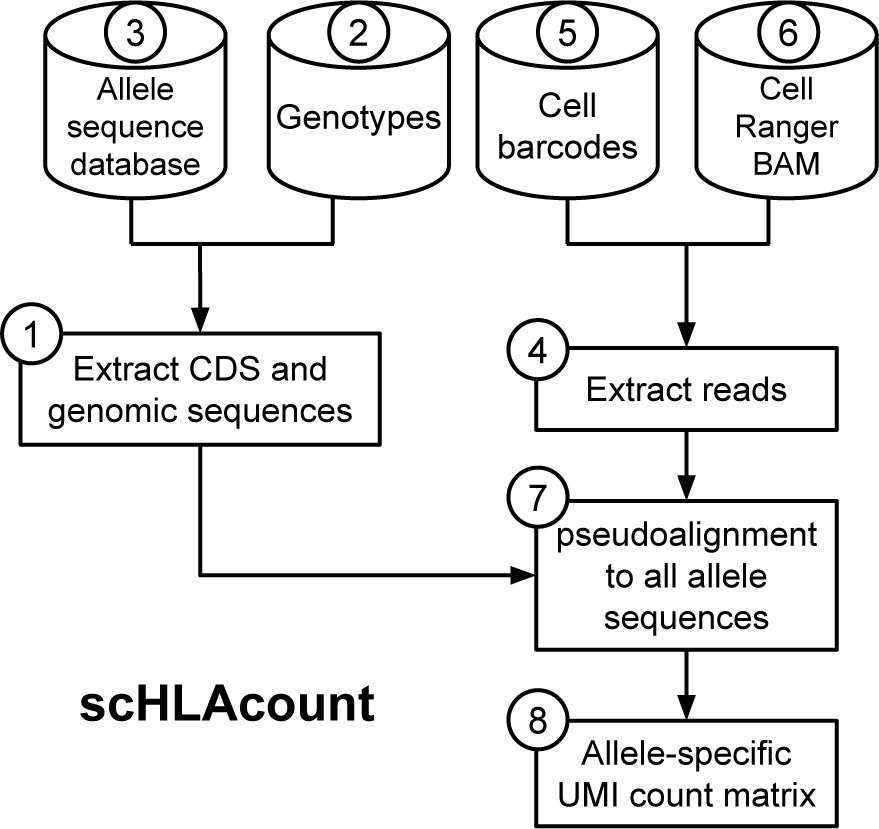
scHLAcount pipeline illustration.

We demonstrate that HLA genes present cell-type specific expression (Boegel et al. 2018) and that HLA loss of expression can be evaluated per-cell and per-cluster. Using five AML samples published in (Petti et al. 2019) for which HLA class I and class II genotypes were provided by the authors, we demonstrate the ability to find cell type specific allele bias when cell types have been annotated using marker genes. We also analyze data from two Merkel cell carcinoma (MCC) patients published in (Yost et al. 2019) and extend their finding that HLA class I expression is lost, to show that this expression loss may be allele-specific. Both datasets illustrate that most molecules in 5’ GEX data can be assigned to a specific allele when the individual is heterozygous, resulting in dataset-wide estimates of allele bias in molecule counts.

### Acute myeloid leukemia (AML)

10x Genomics Chromium 5’ GEX library data derived from five subjects with AML, as described in (Petti et al. 2019) was reanalyzed. Genotypes for HLA-A, -B, -C, -DRB1, and -DQB1 at two-field resolution were provided by the authors.

Given the genotypes, we built custom diploid references; the allele from GRCh38 primary assembly was used for genes HLA-DPA1, DPB1, and DQA1 for which genotypes were not available. Raw scHLAcount molecule counts are summarized in Table S2. Molecule counts were normalized with the following formula:

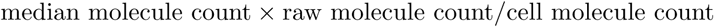

median molecule count × raw molecule count/cell molecule count Normalization and dimensionality reduction of the gene expression matrix generated by Cell Ranger v2.1.1 was performed using Seurat v3.0.2 (Stuart et al. 2019). For all the biallelic genes in each subject, we calculated the average normalized expression per gene and the fraction of the normalized expression for each allele of the nine cell types with at least 100 cells assigned. As observed in the T cell dataset, some genes had more expression of one allele than the other. Results for subject 809653 with the class II gene HLA-DRB1 are listed in Table 1 and visualized on a t-SNE dimensionality reduction plot in Figure 2a,b. Depending on cell type, we observe 42% to 54% allelic bias for the DRB*01:03 allele. This allele preference does not show a trend with average expression. For the same subject, we also observe a 27% to 41% allelic bias for C*07:02 depending on cell type (Figure 2c,d; Table 2).

**Table 1:** Normalized expression and allele-specific expression of HLA-DRB1 for subject 809653 from (Petti et al. 2019), stratified by cell type. Average is taken over all cells assigned to a particular cell type.

| Cell type | # cells | % of DRB1 molecules assigned to 01:03 allele | Avg. HLA-DRB1 normalized expression |
| --- | --- | --- | --- |
| ERY | 3,728 | 41.9 | 0.238 |
| T-CELL | 10,942 | 44.8 | 0.741 |
| PRE-B-CELL | 336 | 47.4 | 1.162 |
| B-CELL | 868 | 47.4 | 14.185 |
| HSC | 2,261 | 52.1 | 5.247 |
| MEP | 560 | 53.0 | 3.411 |
| DEND (M) | 620 | 53.7 | 17.602 |
| ERY (CD34+) | 432 | 53.9 | 2.153 |
| MONO | 1,366 | 54.0 | 7.390 |

**Table 2:** Normalized expression and allele-specific expression of HLA-C for subject 809653 from (Petti et al. 2019), stratified by cell type. Average is taken over all cells assigned to a particular cell type.

| Cell type | # cells | % of HLA-C molecules assigned to 07:02 allele | Avg. HLA-C normalized expression |
| --- | --- | --- | --- |
| B-CELL | 868 | 26.7 | 5.184 |
| MONO | 1,366 | 32.7 | 5.813 |
| PRE-B-CELL | 336 | 33.9 | 3.266 |
| DEND (M) | 620 | 35.1 | 3.890 |
| T-CELL | 10,942 | 37.0 | 8.926 |
| HSC | 2,261 | 38.8 | 3.281 |
| MEP | 560 | 40.3 | 2.578 |
| ERY (CD34+) | 432 | 40.9 | 2.429 |
| ERY | 3,728 | 41.0 | 0.386 |

**Figure 2:**
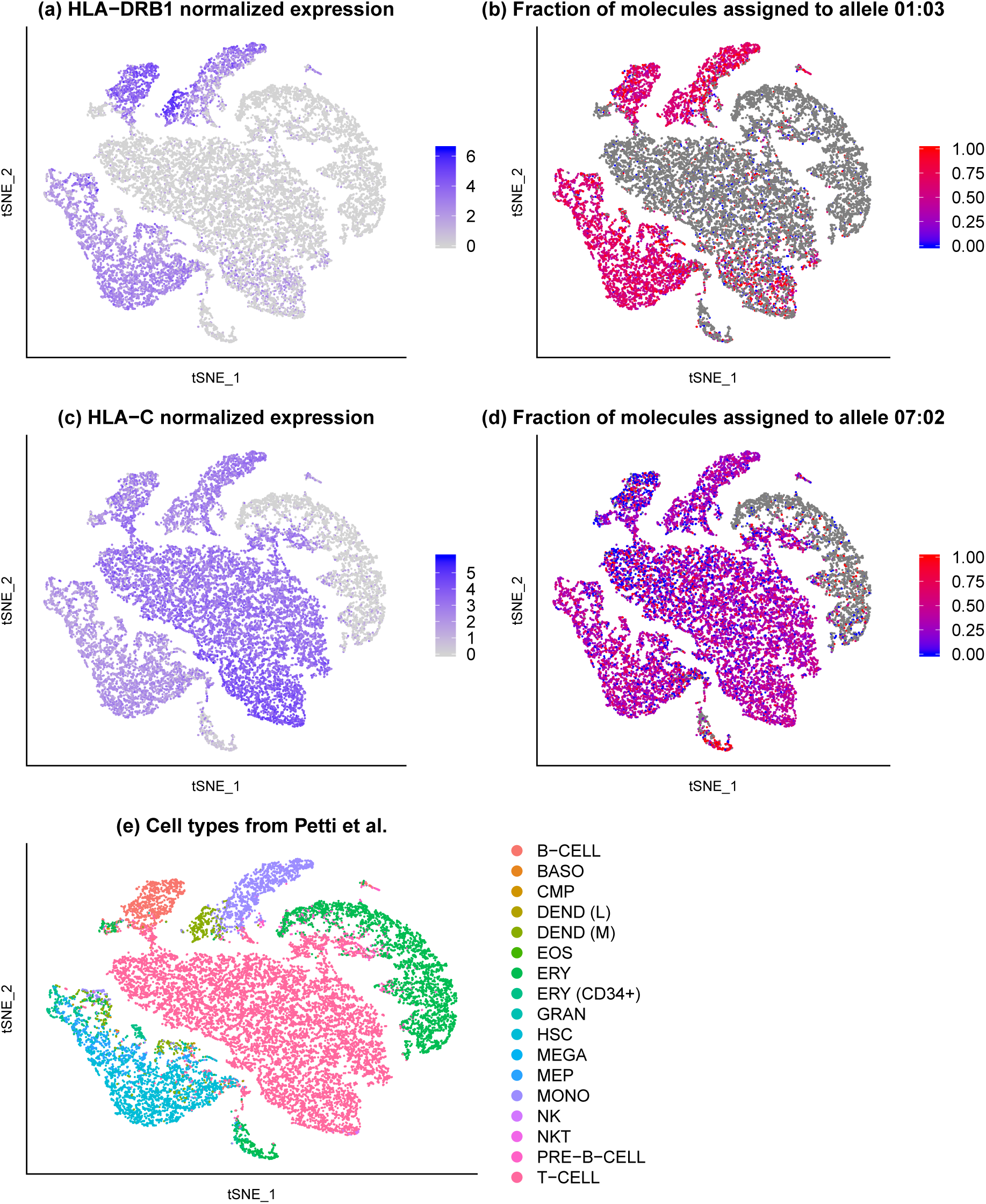
(a) For each cell, color indicates log_2_(1 + normalized expression) of HLA-DRB1. (b) For each cell, color indicates the fraction of HLA-DRB1 molecules assigned to an allele that are assigned to the 01:03 allele of subject 809653. Overall, 95.4% of HLA-DRB1 molecules are assigned to an allele. Gray cells have no HLA-DRB1 molecules assigned to an allele. (c) log_2_(1 + normalized expression) of HLA-C (d) (e) Cell types as inferred in (Petti et al. 2019).

### Merkel cell carcinoma (MCC)

Genotypes for genes HLA-A, -B, and -C for the discovery and validation subjects in (Paulson et al. 2018) were provided by the authors. Here, alleles not explicitly reported in their publication are given a placeholder name (e.g. ‘A1’). Using scHLAcount with a custom reference for the diploid genotype of genes HLA-A, -B, and -C (and GRCh38 primary assembly alleles for the class II genes) we calculated allele-resolved molecule counts. Raw molecule counts were normalized as described above.

Normalization, dimensionality reduction, and clustering was performed using Seurat v3.0.2 (Stuart et al. 2019) following Paulson et al (Paulson et al. 2018). For the discovery subject, we used the filtered expression matrices for tumor and PBMC samples available at GEO accession GSE117988; for the validation subject, the matrix is available at GSE118056.

### Discovery subject

For this subject, the “tumor dataset” comprises cells taken from two time points in treatment; the “PBMC dataset” comprises cells taken from four time points in treatment. Unsupervised clustering of the tumor dataset resulted in 15 clusters. As described in Paulson et al (Paulson et al. 2018), we identified 11 of these clusters comprising 7,131 cells as putative tumor cells using the tumor marker genes NCAM1, KRT20, CHGA, and ENO2 and the non-tumor marker genes CD3D, CD34, CD61, and Fibronectin. The remaining four clusters contained 300 putative normal cells.

Due to the nature of 3’ GEX data, nearly all reads are sequenced from the opposite end of the HLA-A transcript from the variable sites used to define HLA types S1. These variable sites are mostly located in exons 2 and 3, while the 3’ end of the transcripts are mostly homologous between the class I genes Boegel et al. 2018. As a result of the coverage distribution of 3’ GEX data, very few HLA-A molecules could be assigned to an allele. We observe far fewer molecules from scHLAcount compared to Cell Ranger, especially in gene HLA-B because UMIs that only contain reads from the 3’ end of the transcript will be ambiguously aligned to all class I genes and the molecule will not be counted by our algorithm.

As previously reported, HLA-B expression is markedly less in the tumor compared to non-tumor cells and PBMC (Table 4). Additionally, HLA-A and HLA-C expression appears to be reduced in tumor cells.

**Table 3:** scHLAcount analysis of discovery patient tumor (2 time points) and PBMC (4 time points) and validation patient tumor and PBMC (1 time point each) (Paulson et al. 2018). Raw molecule counts for genes A, B, and C are compared to Cell Ranger counts normalized to 1.0. GEX = gene expression

| Subject | Assay type | Custom diploid reference | % molecules assigned to an allele | Custom diploid reference | % molecules assigned to an allele | Custom diploid reference | % molecules assigned to an allele |
| --- | --- | --- | --- | --- | --- | --- | --- |
|  |  | HLA-A |  | HLA-B |  | HLA-C |  |
| Discovery (Tumor) | 3' GEX | 0.866 | 5.34 | 0.391 | 40.76 | 0.639 | 64.31 |
| Discovery (PBMC) | 3' GEX | 0.855 | 6.42 | 0.449 | 45.98 | 0.767 | 67.94 |
| Validation (Tumor) | 5' GEX | 0.878 | 81.17 | 0.896 | 91.69 | 0.745 | 80.68 |
| Validation (PBMC) | 5' GEX | 1.050 | 87.71 | 1.073 | 94.41 | 1.033 | 89.65 |

**Table 4:** Average overall and allele-specific expression of HLA class I genes in the discovery subject of (Paulson et al. 2018).

| Gene<br>Genotype | Tumor cells<br>(n=7,131) |  | Non-tumor cells<br>(n=300) |  | PBMC<br>(n=12,874) |  |
| --- | --- | --- | --- | --- | --- | --- |
|  | Average<br>normalized<br>expression | % molecules<br>assigned to<br>allele 1 | Average<br>normalized<br>expression | % molecules<br>assigned to<br>allele 1 | Average<br>normalized<br>expression | % molecules<br>assigned to<br>allele 1 |
| HLA-A<br>A1/A2 | 0.724 | 24.98 | 3.392 | 43.78 | 1.958 | 40.83 |
| HLA-B<br>35:02/B2 | 0.115 | 76.11 | 3.156 | 61.70 | 1.713 | 63.97 |
| HLA-C<br>C1/C2 | 0.209 | 49.54 | 3.802 | 59.58 | 1.918 | 59.17 |

### Validation subject

For this subject, the “tumor dataset” and “PBMC dataset” comprise cells taken from a single time point after relapse. Unsupervised clustering of all cells together resulted in 18 clusters. As described in Paulson et al (Paulson et al. 2018), we identified seven of these clusters comprising 4,682 cells as putative tumor cells using the tumor marker genes NCAM1, KRT20, Large T Antigen, and Small T Antigen. Only 17 of these cells originated from the PBMC dataset. The remaining 6,209 cells were designated putative normal cells and comprised 5,731 cells from the PBMC dataset and 478 cells from the tumor dataset, which Paulson et al. 2018 identified as tumor-infiltrating leukocytes and tumor-associated macrophages (Figure 2e).

Compared to Cell Ranger molecule counts, we inferred more molecules for the PBMC dataset and fewer molecules for the tumor dataset. At least 80% of scHLAcount molecules were assigned to an allele for class I genes (Table 3).

Dividing cells into tumor and normal as described above, we corroborate the observation from Paulson et al. 2018 that HLA-A expression is greatly reduced in tumor cells compared to infiltrating immune cells (Figure 3a). No marked allele-specific bias in expression is observed in cells in either category. Additionally, we observe decreased expression of HLA-B and HLA-C in tumor cells (Figure 3c,e). While non-tumor cells display approximately balanced expression of the two alleles of these genes, tumor cells have only 13% of allele-resolved HLA-B expression from allele 35:01 and 6% of allele-resolved HLA-C expression from allele ‘C1’ (Table 5).

**Table 5:** Average overall and allele-specific expression of HLA class I genes in the validation subject of (Paulson et al. 2018).

| Gene<br>Genotype | Tumor cells<br>(n=4862) |  | Non-tumor cells<br>(n=6209) |  |
| --- | --- | --- | --- | --- |
|  | Average<br>normalized<br>expression | % molecules<br>assigned to<br>allele 1 | Average<br>normalized<br>expression | % molecules<br>assigned to<br>allele 1 |
| HLA-A<br>02:01/A2 | 0.060 | 39.7 | 4.154 | 56.8 |
| HLA-B<br>35:01/B2 | 0.511 | 13.4 | 5.172 | 50.4 |
| HLA-C<br>C1/C2 | 0.327 | 6.3 | 4.991 | 46.8 |

**Figure 3:**
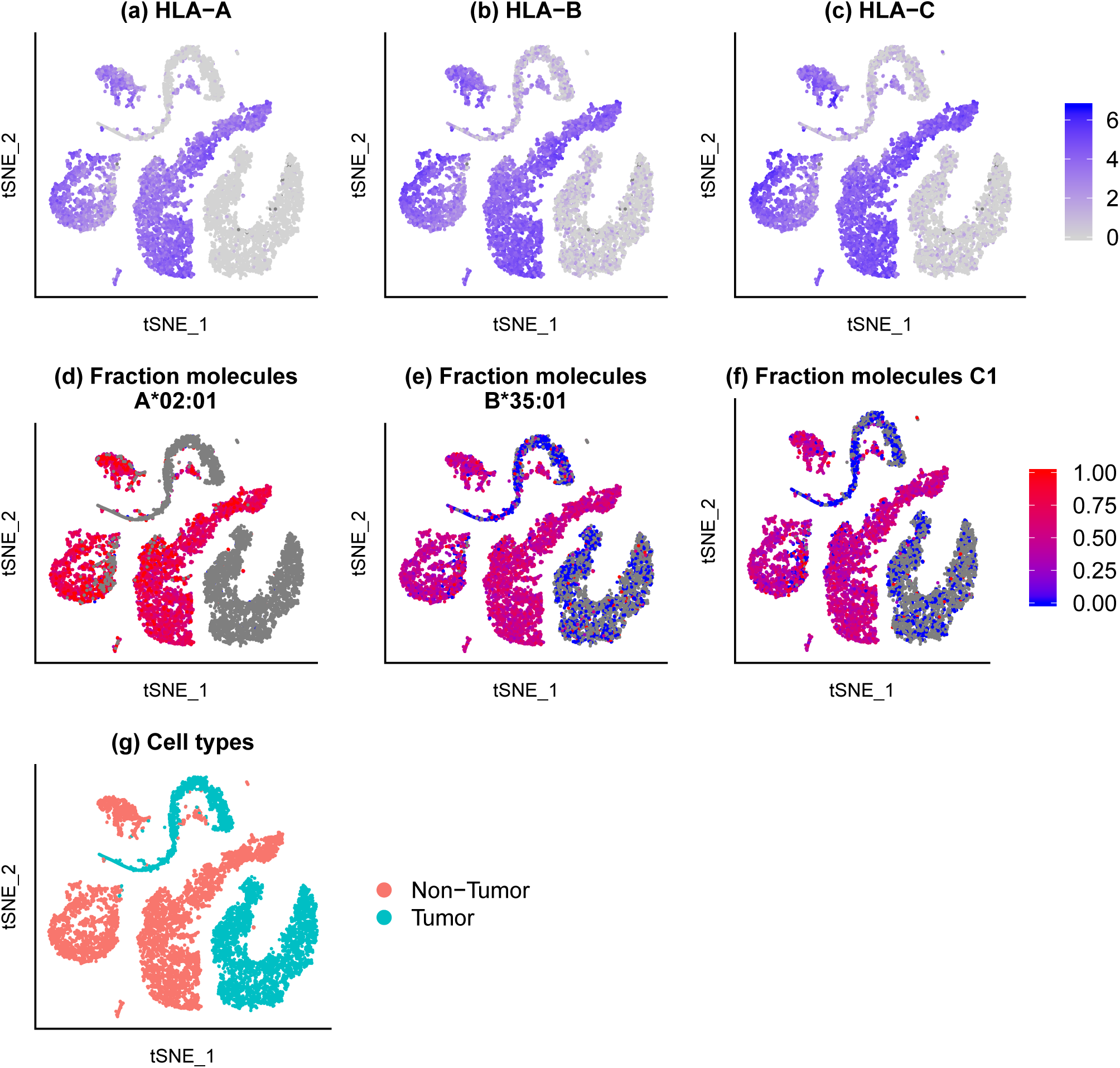
log_2_(1 + normalized expression) of HLA-A (a) HLA-B (b) and HLA-C (c) and allele preference for HLA-A*02:01 (d) HLA-B*35:01 (e) and HLA-C1 (f) for the validation subject of (Paulson et al. 2018). Values are plotted per cell; aggregate statistics shown in Table 5. (g) Cell types inferred using marker genes.

## Discussion

Tumor evasion of immunotherapy is of growing concern, as novel and expensive treatment modalities find themselves stymied by this evolutionary response. scHLAcount provides a simple way to assign reads from scRNA-seq experiments to MHC alleles. This is a powerful tool for investigating allele-specific expression, loss of heterozygosity, and mutational or epigenetic suppression of HLA expression in tumor immune-evasion. Additionally, using a personalized reference and counting with scHLAcount often recovers more molecules than using the standard reference and counting with Cell Ranger. This has the potential to improve gene expression based clustering in cells where MHC genes are a major component of the expression profile. scHLAcount could be extended to also apply to any other locus where there is common structural variation present in the human population. The approach of using De Bruijn graphs to improve isoform and haplotype quantification has been considered before (Patro et al. 2017; Bray et al. 2016), but has not yet been applied to scRNA-seq data until this study. A recent pre-print (Tian et al. 2019) genotyped individual cells for HLA class I using scRNA-seq data but did not address allele-specific expression on the molecule level.

We have found that 5’ GEX data is preferable to 3’ GEX data for genotyping and assigning molecules to alleles, because the sequencing coverage is not as limited to one end of the transcript (Figure S1). Since the three class I genes have considerable sequence homology except in exons 2 and 3 and virtually all of the coverage of 3’ GEX data falls in the later exons, few UMIs have a read from the variable exons and could be assigned to a specific allele, and many UMIs have reads only in regions homologous among the 3 genes and thus these molecules are not counted.

## Methods and Materials

The numbered steps in Figure 1 correspond to the numbers in parentheses in this section.

To make FASTA files of the coding and genomic sequences of the alleles present in the sample (1), users need to provide a list of genotypes (2) and download the IMGT/HLA allele sequence database (3). These genotypes can be assayed by specialized molecular tests, such as sequence-specific oligonucleotide probe PCR (PCR-SSOP), sequence-specific primed PCR (PCR-SSP), or Sanger sequence-based typing (SBT) (Erlich 2012). Alternatively, algorithms for sequence-based typing from next-generation sequencing reads of the genome, exome, or transcriptome utilize comprehensive allele databases such as IMGT/HLA (Robinson et al. 2015) to successfully infer genotypes (reviewed in Bauer et al. 2018). Following the pseudoalignment approachBray et al. 2016, scHLAcount builds two colored De Bruijn graph indexes, one containing the CDS sequences and one containing genomic sequences, using a k-mer length of 24.

Reads aligned to the MHC region (GRCh38 coordinates chr6:28510120-33480577) (4) corresponding to valid cell barcodes (5) are extracted from the BAM file (6). Each read is first pseudo-aligned to the CDS graph, yielding the set of alleles that could have generated the read (also referred to as the equivalence class) Bray et al. 2016. If there is no alignment of at least 60 bases (2 mismatches are permitted outside the initial seed kmer), the read is pseudo-aligned to the genomic sequence graph and retained if the alignment is at least 60 bases. (7) In the datasets studied, less than 5% of reads that failed to align to the CDS were successfully aligned to the genomic sequence. This genomic alignment step is intended to rescue reads that may be haplotype specific in 3’ or 5’ UTR regions. It also provides a mechanism to handle single nuclei RNA-seq libraries.

Reads sharing a cell barcode and unique molecular identifier (UMI) are assumed to originate from the same RNA molecule. At recommended sequencing depths with modest sequence saturation, there are typically 1-3 reads per UMI. Individual reads may have different equivalence classes according to their pseudoalignment. We ignore reads whose equivalence class contains more than one gene. If more than half of the reads from a molecule are assigned to a particular gene, that molecule will get counted to one of its alleles (e.g. HLA-A 02:01), based on the constituent reads’ equivalence classes. In the case of ambiguity, it will get counted to that gene (e.g. HLA-A) instead. The output is a sparse molecule count matrix (8) where each column corresponds to a barcode in the provided cell barcode list, and each row corresponds to an allele.

## Acknowledgements

We thank Kelly Paulson and Allegra Petti for providing HLA genotypes of the subjects in the MCC and AML studies respectively.

## Author Contributions

Conceptualization, A.M.B. and I.T.F.; Methodology, C.A.D.; Software, C.A.D. and I.T.F.; Investigation, C.A.D.; Data Curation, C.A.D. and I.T.F.; Writing - Original Draft, C.A.D. and I.T.F.; Writing - Review & Editing C.A.D., M.J.T.S., P.J.M., A.M.B. and I.T.F.; Visualization, C.A.D.; Supervision, A.M.B. and P.J.M.

## Supplementary Material

### GRCh38 primary assembly alleles

Genotypes present in GRCh38 primary assembly were inferred using Kourami v0.9.6 (H. Lee and Kingsford 2018). 2 million 200bp error-free reads were simulated from GRCh38 Chr6:28510120-33480577, which is approximately 80-fold coverage of the region. Reads were aligned to the Kourami reference panel and genotypes were inferred; all listed genotypes had 100% sequence identity with respect to the corresponding database sequence.

A*03:01:01G B*07:02:01G

C*07:02:01G

DQA1*01:02:01G

DQB1*06:02:01G

DRB1*15:01:01G

DPA1*01:03:01G

DPB1*04:01:01G

### Computational performance

On the scRNA-seq dataset from donor 4 from 10X Genomics 2019, scHLAcount analyzed 58 million reads aligned to the MHC region in 83 minutes (55 minutes spent genotyping; 28 minutes spent counting). Maximum memory usage was 1.5 GB.

**Table S1:** Genotypes for subjects from Petti et al. 2019, provided by the authors in personal communication with permission to include here. Genotypes shared with the GRCh38 primary assembly are in **bold text**.

| Subject | HLA-A | HLA-B | HLA-C | HLA-DQB1 | HLA-DRB1 |
| --- | --- | --- | --- | --- | --- |
| 508084 | 68:01/01:01 | **07:02**/27:05 | **07:02**/07:04 | 05:01/**06:02** | 01:01/**15:01** |
| 548327 | 68:01/02:06 | 51:01/44:05 | 02:02 | 02:02/02:01 | 07:01/03:01 |
| 721214 | **03:01**/01:01 | 18:01/14:01 | 08:02/07:40 | 03:02/02:02 | 07:01/04:03 |
| 782328 | 32:01 | 37:01/15:01 | 06:02/03:04 | 05:01/03:02 | 04:01/10:01 |
| 809653 | 68:02/31:01 | 27:05/14:02 | 08:02/**07:02** | 03:01 | 11:01/01:03 |

**Table S2:** Using the custom diploid reference or GRCh38 allele as denoted, raw molecule count for each gene is compared to Cell Ranger counts normalized to 1.0. Subject 548327 is homozygous for HLA-C, Subject 782328 is homozygous for HLA-A, and Subject 809653 is homozygous for HLA-DQB1.

| Subject | Custom<br>diploid<br>reference | % molecules<br>assigned to<br>an allele | Custom<br>diploid<br>reference | % molecules<br>assigned to<br>an allele | Custom<br>diploid<br>reference | % molecules<br>assigned to<br>an allele | Custom<br>diploid<br>reference | % molecules<br>assigned to<br>an allele |
| --- | --- | --- | --- | --- | --- | --- | --- | --- |
|  | HLA-A |  | HLA-B |  | HLA-C |  | HLA-DQB1 |  |
| 508084 | 1.039 | 95.13 | 1.066 | 87.22 | 0.885 | 60.77 | 1.028 | 95.89 |
| 548327 | 1.165 | 86.26 | 1.061 | 93.09 | 1.032 | n/a | 2.721 | 2.27 |
| 721214 | 1.180 | 69.44 | 1.137 | 90.09 | 0.908 | 93.63 | 3.319 | 98.95 |
| 782328 | 1.154 | n/a | 0.880 | 63.95 | 0.957 | 89.77 | 1.010 | 99.15 |
| 809653 | 1.083 | 87.21 | 1.154 | 96.53 | 0.911 | 91.74 | 1.070 | n/a |
| Subject | Custom<br>diploid<br>reference | % molecules<br>assigned to<br>an allele | GRCh38<br>allele | GRCh38<br>allele | GRCh38<br>allele |  |  |  |
|  | HLA-DRB1 |  | HLA-DPA1 | HLA-DPB1 | HLA-DQA1 |  |  |  |
| 508084 | 1.641 | 74.60 | 1.135 | 1.024 | 1.086 |  |  |  |
| 548327 | 1.920 | 89.52 | 1.180 | 1.172 | 2.087 |  |  |  |
| 721214 | 1.745 | 89.05 | 1.217 | 1.050 | 2.058 |  |  |  |
| 782328 | 1.125 | 92.12 | 1.276 | 1.078 | 1.274 |  |  |  |
| 809653 | 1.066 | 95.43 | 1.136 | 1.050 | 1.455 |  |  |  |

**Figure S1:**
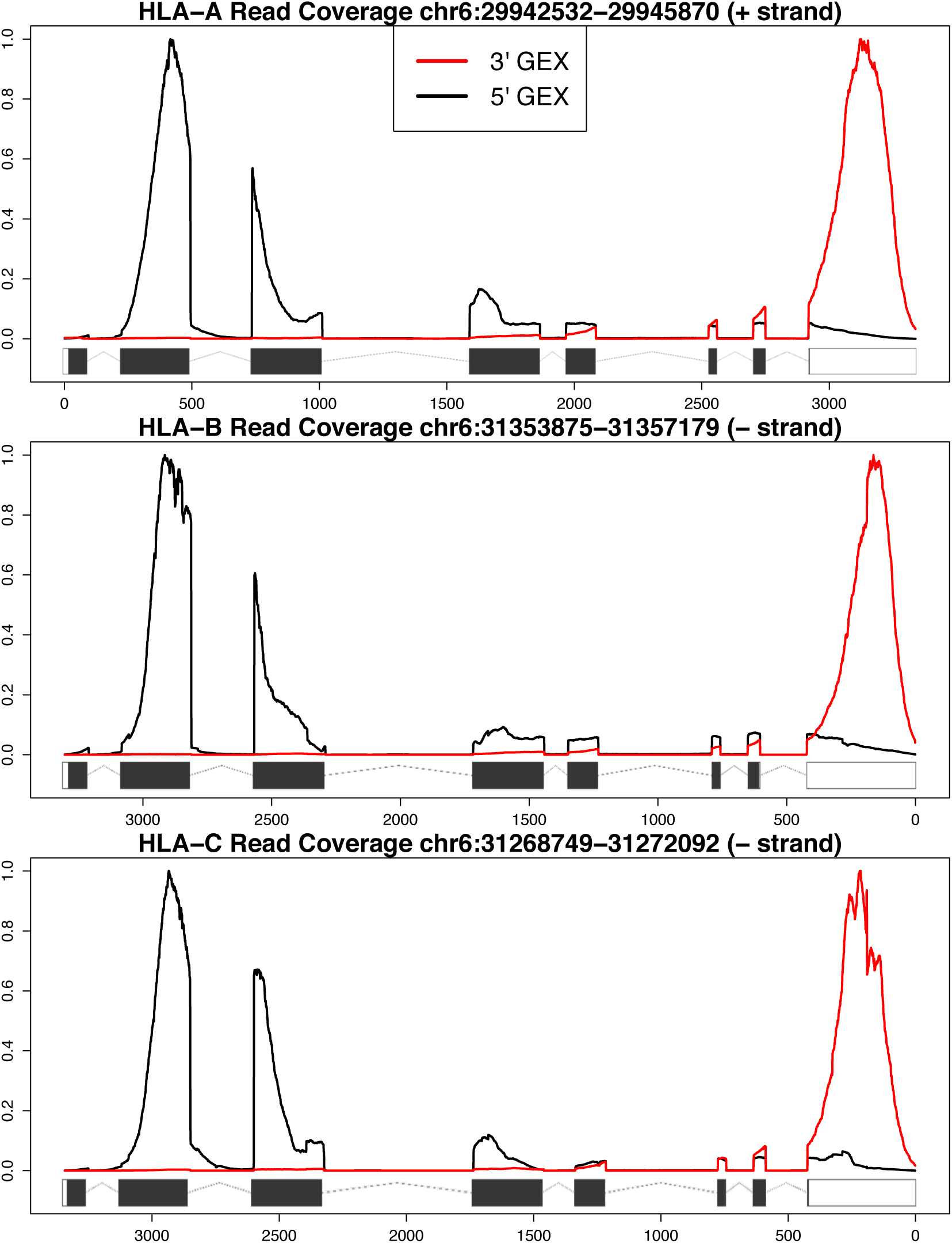
Read coverage of HLA Class I genes for 3’ GEX and 5’ GEX. Minimum and maximum coverage for each assay in the region shown is normalized to 0 and 1 respectively. The 3’ dataset is merged from SRR7722937-SRR7722942 and the 5’ dataset is SRR7692286 (Paulson et al. 2018). GEX = gene expression

